# Use-dependent learning biases the initial state but not the learning dynamics of implicit adaptation

**DOI:** 10.64898/2025.12.12.694069

**Authors:** Yuxuan Luo, Kunlin Wei

## Abstract

Motor adaptation arises from multiple learning mechanisms, including use-dependent learning (UDL) driven by repetition and implicit error-based learning (EBL) driven by motor prediction errors. Although both mechanisms contribute implicitly to shaping movement execution, whether these two mechanisms interact remains unclear. The present study used an error-clamp (EC) task to isolate implicit EBL and directly examined whether UDL affects subsequent implicit adaptation. Participants first performed extensive single-target repetitive reaches to induce use-dependent biases and then immediately transitioned to an EC task. We manipulated whether the UDL bias was aligned with, opposed to, or neutral toward the to-be-adapted direction in three separate groups of participants. Results demonstrate that UDL robustly shifted the initial state of adaptation, but did not alter the learning dynamics: all groups showed comparable trial-by-trial adaptation and converged to similar asymptotic levels despite their different initial states. These findings support an independent rather than an interactive relationship between UDL and implicit EBL, highlighting that execution-level motor learning components are additive.

## Introduction

The human motor system continuously adapts to environmental perturbations and bodily changes to maintain accurate performance. This adaptive capacity emerges from multiple learning mechanisms: reinforcement learning (RL), use-dependent learning (UDL), and error-based learning (EBL; Wolpert et al., 2011). Each mechanism relies on distinct information sources as learning signals. RL optimizes motor actions for rewarding outcomes through exploration and reinforcement (Sutton & Barto, 1998). UDL biases future movements toward recently executed patterns through repetition (Diedrichsen et al., 2010; Tsay et al., 2022). EBL recalibrates motor output based on motor errors, with the predominant error signal being prediction error—the discrepancy between intended and actual movement feedback (Shadmehr et al., 2010; Zhang et al., 2024).

These learning mechanisms operate across different levels of the motor control hierarchy. Within the framework proposed by Wolpert and colleagues (2011), motor learning involves information extraction, decision making, and control processes. RL primarily engages decision-making through exploration and action selection based on outcomes. EBL encompasses both explicit strategies accessible to conscious awareness and implicit recalibration that automatically adjusts internal models (McDougle et al., 2015). UDL represents the most automatic form of learning, operating purely at the execution level through history-dependent changes in sensorimotor representations (Tsay et al., 2022).

Despite growing evidence that these mechanisms coexist during motor adaptation (e.g., Haith & Krakauer, 2013; Spampinato & Celnik, 2021), their interactions remain poorly understood. Recent work has focused primarily on the interaction between RL and EBL, revealing that reward signals can regulate both learning rate and memory retention of EBL through metacognitive processes (Sugiyama et al., 2023). For instance, punishment accelerates error-based adaptation while reward enhances its retention (Galea et al., 2015), and task success attenuates implicit recalibration (Kim et al., 2019). These findings suggest that high-level processes related to decision and action selection can gate low-level implicit learning based on motor errors.

However, interactions between implicit, execution-level mechanisms remain largely unexplored. Specifically, whether and how UDL and implicit EBL interact has not been systematically investigated. This gap is surprising given their overlapping neural substrates: UDL involves Hebbian-like plasticity in primary motor cortex (M1) and premotor areas (Classen et al., 1998; Verstynen & Sabes, 2011), while implicit EBL relies on cerebellar-M1 loops for sensorimotor recalibration (Galea et al., 2011; Schlerf et al., 2012). Both mechanisms thus converge on M1 as the final common pathway for motor execution. The only related study on the relationship of UDL and EBL was conducted by Diedrichsen and colleagues (2010), who found that UDL induced motor biases that constituted initial movement errors that could be corrected in subsequent movement attempts. However, whether and how UDL and implicit EBL interact remains an open question. Understanding their potential interactions carries both theoretical importance and practical implications. For instance, current rehabilitation and skill training protocols often combine repetitive practice with error-based training (e.g., Marchal-Crespo et al., 2017; Reinkensmeyer et al., 2016). If use-dependent biases modulate error sensitivity, these training protocols could be optimized accordingly.

The present study directly examines whether UDL, the repetition-induced motor bias, affects subsequent implicit EBL. Our participants first performed extensive repetitive reaches to a fixed target to establish a UDL bias, then immediately transitioned to a single-target error-clamp (EC) adaptation task that primarily engages implicit EBL (Morehead et al., 2017). Previous research indicates that UDL bias “attracts” subsequent reaches intended for nearby targets toward the repeated direction (e.g., Huang et al., 2011; Marinovic et al., 2017; Tsay et al., 2022; Verstynen & Sabes, 2011), potentially influencing the subsequent EC task. Thus, the interaction hypothesis proposes that UDL-induced biases could alter the EBL process itself, showing as a directional attraction that modulates the rate or asymptotic extent of subsequent implicit adaptation. In contrast, the independent hypothesis posits that UDL and EBL operate independently, with the adaptive process following UDL remaining unchanged.

To distinguish between independent and interaction hypotheses, we manipulated the spatial relationship between UDL attractors and the subsequent EC target direction using three experimental conditions. In the Attracting condition, the UDL bias was aligned with the direction of the subsequent implicit adaptation. In the Repelling condition, the UDL bias directly opposed this to-be-adapted direction. In the Control condition, UDL training occurred at the EC target location, establishing a baseline without directional bias relative to the subsequent adaptation. This design yields distinct predictions for each hypothesis. Under the independent hypothesis, repetition-induced biases would merely provide different starting conditions for EBL, leaving error-based adaptation rates and extents unchanged across conditions.Alternatively, under the interaction hypothesis, we expect systematic differences in learning dynamics depending on the pattern of pre-existing UDL biases. Specifically, compared to the Control condition, EBL might proceed more rapidly in the Attracting condition and more slowly in the Repelling condition.

## Method

### Participants

Sixty right-handed students from Peking University participated in this experiment (mean age: 22.16 ± 3.16 years; 16 males, 45 females). All participants reported normal or corrected-to-normal vision and no history of neurological or motor impairments. Participants were naive to the experimental hypotheses and had no prior experience with similar motor learning paradigms. All participants provided written informed consent before participation and received monetary compensation or course credit upon completion.

### Apparatus and Stimuli

Participants performed center-out reaching movements using a Wacom Intuos Pro digitizing tablet (PTK-1240; active area: 487.7 × 304.8 mm; sampling rate: 125 Hz) with a wireless stylus. Visual stimuli were displayed on a monitor (509.2 × 286.4 mm; 1920 × 1080 pixels; 60 Hz refresh rate) positioned 196 mm directly above the tablet surface. Participants were seated comfortably with the tablet positioned at approximately waist height. An opaque acrylic screen and wooden enclosure completely occluded direct vision of the hand and stylus, ensuring that participants relied solely on visual cursor feedback.

Visual stimuli consisted of a white circular cursor (3 mm diameter), a yellow starting circle (5 mm diameter), and pink circular targets (4 mm diameter). All targets appeared at a fixed radial distance of 80 mm from the start position. The background was black to maximize contrast. Stimuli were programmed in MATLAB R2021a using Psychtoolbox-3.

### Experimental Design

Each trial followed a standardized sequence (Figure 1). Participants initiated trials by moving the cursor to the yellow starting position and maintaining position for 0.2 s. A pink target then appeared at one of the designated locations. After a delay period (randomly varied between 0.5-1.0 s), an auditory go-signal (500 Hz tone, 0.1 s duration) prompted movement initiation. This delay ensured adequate preparation time while preventing anticipatory responses. Participants were instructed to make rapid, straight reaching movements toward the target upon hearing the go-signal. Movement amplitude was constrained by terminating trials when cursor displacement exceeded 60 mm from the start position, as indicated by a termination tone. Upon successful target acquisition, the target outline turned green and remained visible for 1.0 s as positive feedback. Cursor feedback was only visible within 15 mm of the start position during return movements to prevent visual guidance during the return to the start.

**Figure 1.**
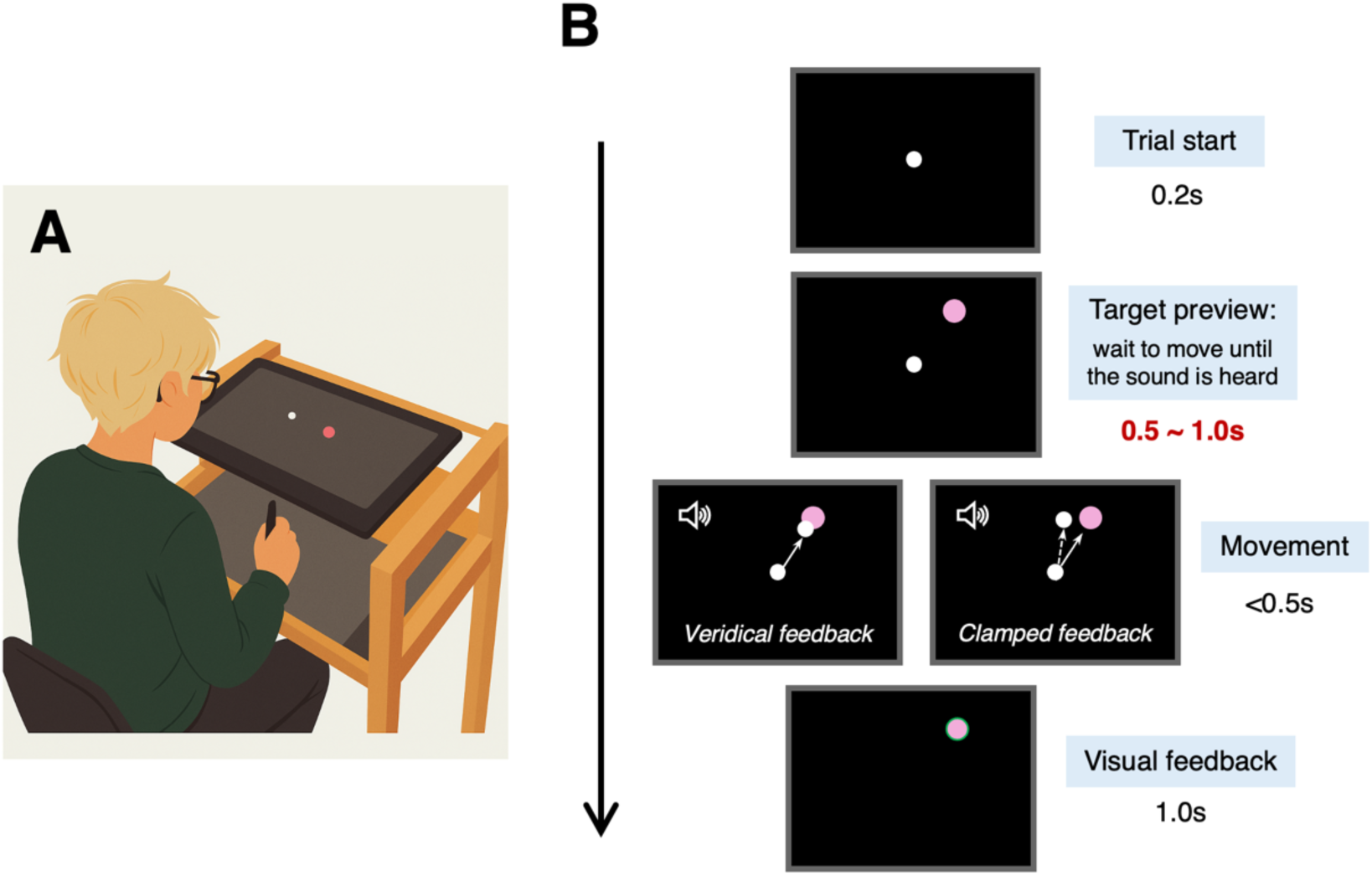
Experimental design. **(A)** Experimental setup. Participants perform reaching movements on a tablet using a stylus held in their right hand. Direct vision of the hand is occluded; participants see only visual stimuli displayed on a monitor positioned above the tablet surface. **(B)** Illustration of a single exemplary trial with veridical or clamped feedback. After the participant’s hand remained at the starting position for 0.2 s, a pink target appears. The participant waits at the starting position for an auditory go-signal, which occurs randomly 0.5-1.0 s after target appearance. In veridical feedback trials, the cursor reflects real-time stylus position. In clamped feedback trials, the cursor is constrained to move along a direction 30° counterclockwise from the target direction. After the hand slices through the target, the target outline turns green (only for successful hits) and remains visible for 1.0 s before the next trial begins.

Three distinct feedback conditions were employed throughout the experiment. During veridical feedback trials, the cursor accurately reflected real-time stylus position throughout the movement, providing natural visual feedback. In no-feedback trials, the cursor disappeared at movement onset, eliminating visual feedback during reaching while preserving target visibility. During EC feedback trials, the cursor trajectory was artificially constrained to move along a fixed angular direction (75° counterclockwise from positive x-axis) regardless of actual hand movement direction or amplitude. Cursor displacement matched actual movement amplitude to maintain realistic feedback characteristics while eliminating directional error information.

We employed a between-subjects design with three experimental groups that differed in the angular relationship between the repetition target and the subsequent EC target (Figure 2). This design allowed us to systematically examine how use-dependent biases with different directional relationships to EC feedback influence subsequent implicit adaptation.

**Figure 2.**
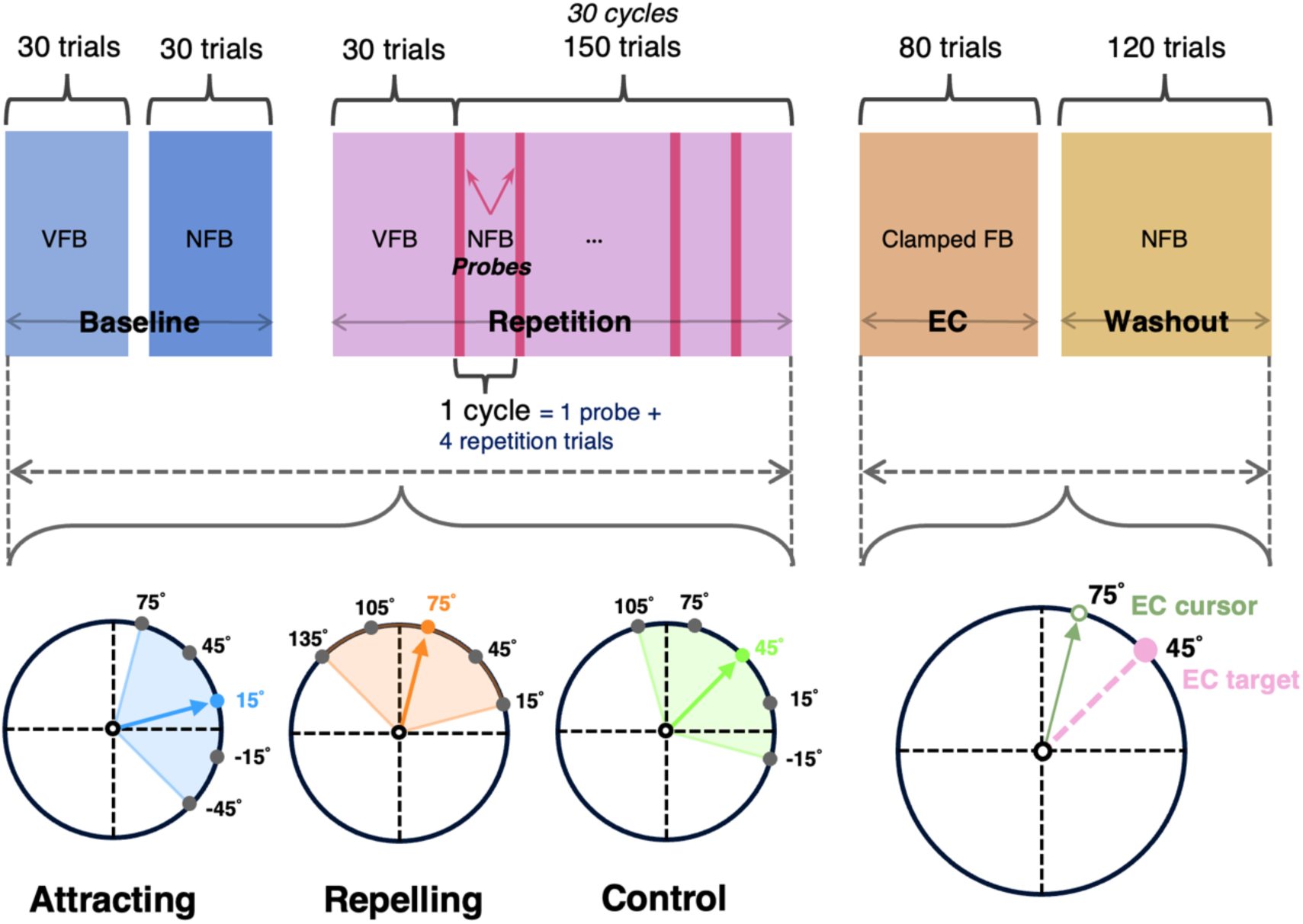
Experimental design. Different colored bars represent distinct experimental blocks. During the baseline and repetition phases, five experimental targets are positioned in different regions of the workspace for the three participant groups, centered at 15°, 45°, and 75°, respectively. The baseline phase consisted of equal number of trials to each of the five targets with or without cursor feedback, to measure the individuals’ inherent directional biases. The repetition phase consisted of 30 repetition trials to the center target, followed by 30 cycles of 5 trials. Each cycle consisted of 4 same repetition trials and one no-feedback probe trial to one of the 5 possible targets. While repetition trials are designed to induce the UDL, probes can assess UDL biases at adjacent directions. Subsequently, all three groups undergo identical implicit adaptation by reaching the same 45° target with clamped feedback. In the final washout phase for measuring learning aftereffect, participants reached the 45° target without cursor feedback.

The Attracting group (n = 20) performed repetitive movements to a target positioned at 15°, creating a 30° clockwise offset from the subsequent 45° EC target. This configuration was designed to induce use-dependent biases that would be congruent with the required counterclockwise adaptation to the 75° clamped cursor. The Repelling group (n = 20) repeated movements to a target at 75°, creating a 30° counterclockwise offset from the EC target. This configuration was designed to induce use-dependent biases that would be incongruent with the required adaptation. The Control group (n = 20) experienced repetition training with the target positioned at 45°, directly aligned with the EC target, allowing assessment of adaptation without directional bias conflicts. For each group, four additional probe targets were positioned symmetrically around the repetition target at ±30° and ±60° intervals to assess the spatial generalization profile of use-dependent effects. All angular positions were measured counterclockwise from the positive x-axis.

All participants completed a standardized practice session before the main experiment. Practice included at least 10 complete cycles (50 trials total) with targets appearing at all five positions in randomized order under veridical feedback conditions. Practice continued until participants demonstrated consistent understanding of task requirements. Subsequently, participants completed 10 no-feedback trials and 10 EC trials to familiarize them with these conditions and ensure they understood the instruction to ignore the cursor during clamped trials.

The main experiment consisted of four sequential phases totaling approximately 30 minutes (Figure 2). During the baseline phase (60 trials), participants completed 30 veridical feedback trials and 30 no-feedback trials with targets appearing randomly and equally likely at all five locations. This phase established individual baseline reaching patterns and inherent directional biases before repetition training. The repetition phase (180 trials) began with 30 veridical feedback trials to the designated repetition target to establish the repeated movement pattern.

Subsequently, participants completed 30 cycles, each containing 4 veridical feedback trials to the repetition target followed by 1 no-feedback trial to a randomly selected probe target; this resulted in 150 trials total. This design maintained repetition frequency while periodically assessing bias at other directions. During the EC phase (80 trials), all participants performed reaching movements to the 45° target with cursor feedback clamped at 75°.This counter-clockwise rotation of cursor feedback would lead to implicit adaptation in the clockwise direction (Morehead et al., 2017). Participants were explicitly instructed that cursor movement was spatially decoupled from their hand movement and to ignore the cursor completely while reaching straight to the target. This instruction emphasized the importance of maintaining target-directed reaching despite misleading visual feedback. Finally, during the washout phase (120 trials), participants completed no-feedback trials exclusively to the 45° target to assess retention and decay of both use-dependent biases and EC adaptation.

### Data Analysis

All data processing and statistical analyses were conducted using MATLAB R2023a and JASP 0.18. Movement direction was quantified as the angular deviation from the target direction at peak velocity, serving as our primary dependent measure. This measure captures the primary direction of the reaching movement while being relatively insensitive to late corrective adjustments. Trials were excluded from analysis based on predetermined criteria to remove outliers and technical artifacts.

These included movement duration exceeding 0.5 s, and trials with apparent equipment malfunctions or participant inattention. Excluded trials comprised 2.47% (Attracting), 1.34% (Repelling), and 1.81% (Control) of total trials. All reported results remained consistent when analyses included these excluded trials. Primary statistical analyses included one-sample t-tests and one-way ANOVAs. For all tests, significance threshold was set at α = 0.05, and data are reported as mean ± SEM.

## Results

To investigate whether UDL influences subsequent implicit EBL, our participants performed a multi-phase reaching task. They first established baseline performance, then underwent extensive repetitive reaching (150 trials to a single repeated target) to induce use-dependent biases, then adapted to EC feedback at a single target positioned at 45°, which engages implicit learning. If UDL biases implicit adaptation, we expected EC learning to proceed more rapidly in the Attracting group and more slowly in the Repelling group compared to the Control group. We focused primarily on directional deviation, quantified as the angular deviation between actual movement direction and target direction, to evaluate baseline performance, and the effects of UDL and implicit EBL.

### Baseline reaching movements exhibit systematic directional biases

Before examining the effects of UDL and EBL, we characterized participants’ inherent movement biases to different targets by analyzing no-feedback trials during the baseline phase. Understanding these baseline biases is crucial for isolating subsequent learning effects from pre-existing tendencies.

Each group of participants reached five targets distributed around their respective repetition target (see Methods). Average reach directions revealed systematic biases with remarkable consistency across groups (Figure 3). Analysis revealed significant counterclockwise biases at 15° (1.21° ± 0.41°, *t(59)* = 2.951, *p* = .005), 105° (3.05° ± 0.76°, *t(39)* = 4.004, *p* < .001), and 135° (2.53° ± 1.09°, *t(19)* = 2.328, *p* = .031). Conversely, significant clockwise biases emerged at 45° (−2.32° ± 0.45°, *t(59)* = -5.104, *p* < .001) and 75° (−4.48° ± 0.49°, *t(59)* = -9.999, *p* < .001).

**Figure 3.**
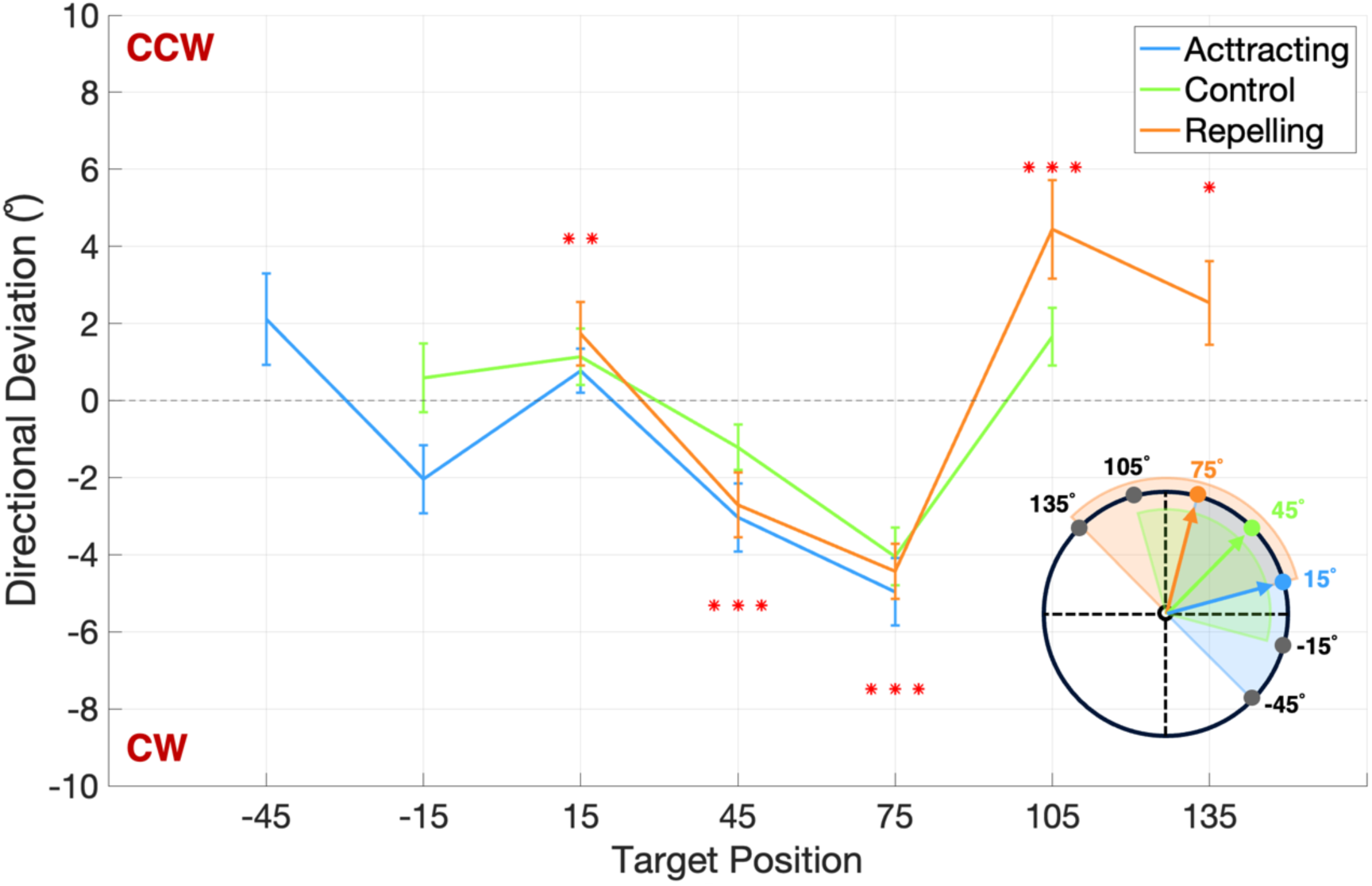
Directional bias measured by baseline trials without visual feedback. Directional biases are shown for different target positions. Each participant group covered a distinct set of targets, thus their directional biases are shown as separate data points with individual error bars. The target sets for each group are illustrated in the inset. Counterclockwise deviations are presented as positive values, and clockwise deviations were presented as negative values. Significant biases against zero are marked with * (p < .05), ** (p < .01), and *** (p < .001). Note that biases were consistent across groups.

Targets at -45° and -15° showed biases of 2.11° ± 1.18° and -0.73° ± 0.66°, respectively, though these did not reach statistical significance, likely due to smaller sample sizes. Notably, the biases at these specific target directions are consistent with a recent large-sample study (Wang et al., 2024). These intrinsic biases provide a baseline for isolating subsequent learning effects from natural movement preferences.

### Repetitive movements induce use-dependent biases toward repeated directions

Following baseline assessment, participants performed extensive repetitive reaching to a single target location (150 trials interspersed with occasional probe trials to other targets). This repetition phase was designed to induce use-dependent biases—a tendency for movements to deviate toward recently repeated directions.

To ensure that any observed biasing effect reflected automatic, execution-level processes rather than conscious strategies, we first verified that participants had adequate preparation time before each movement. Movement preparation times substantially exceeded the required 0.5s (Attracting: 0.96 ± 0.06 s; Control: 1.12 ± 0.04 s; Repelling: 1.16 ± 0.03 s). This extended preparation time minimized the possibility of planning errors, which might manifest as directional deviations (Marinovic et al., 2017; Tsay et al., 2022).

We assessed use-dependent effects by comparing reaching directions during no-feedback probe trials before and after the repetition phase. If UDL exists, it should bias movement direction clockwise when the probe target is counterclockwise to the repetition direction, and counterclockwise when the probe target is clockwise to the repetition direction. Indeed, we observed this UDL effect for most target locations examined (Figure 4). The Attracting group exhibited a significant clockwise deviation at the 75° target (−3.73° ± 0.64°, *F*(1,39) = 8.355, *p* = .006) and a marginally significant counterclockwise deviation at -45° (3.11° ± 0.93°, F(1,39) = 3.568, p = .066) (Figure 4A). Movements to −15° and 45° showed biases of 1.50° ± 0.91° and 1.26° ± 0.81°, respectively, but these did not reach statistical significance. The pattern of significant effects suggests that use-dependent attraction operates most strongly at intermediate angular distances, consistent with the spatial tuning characteristics of UDL reported previously (Marinovic et al., 2017).

**Figure 4.**
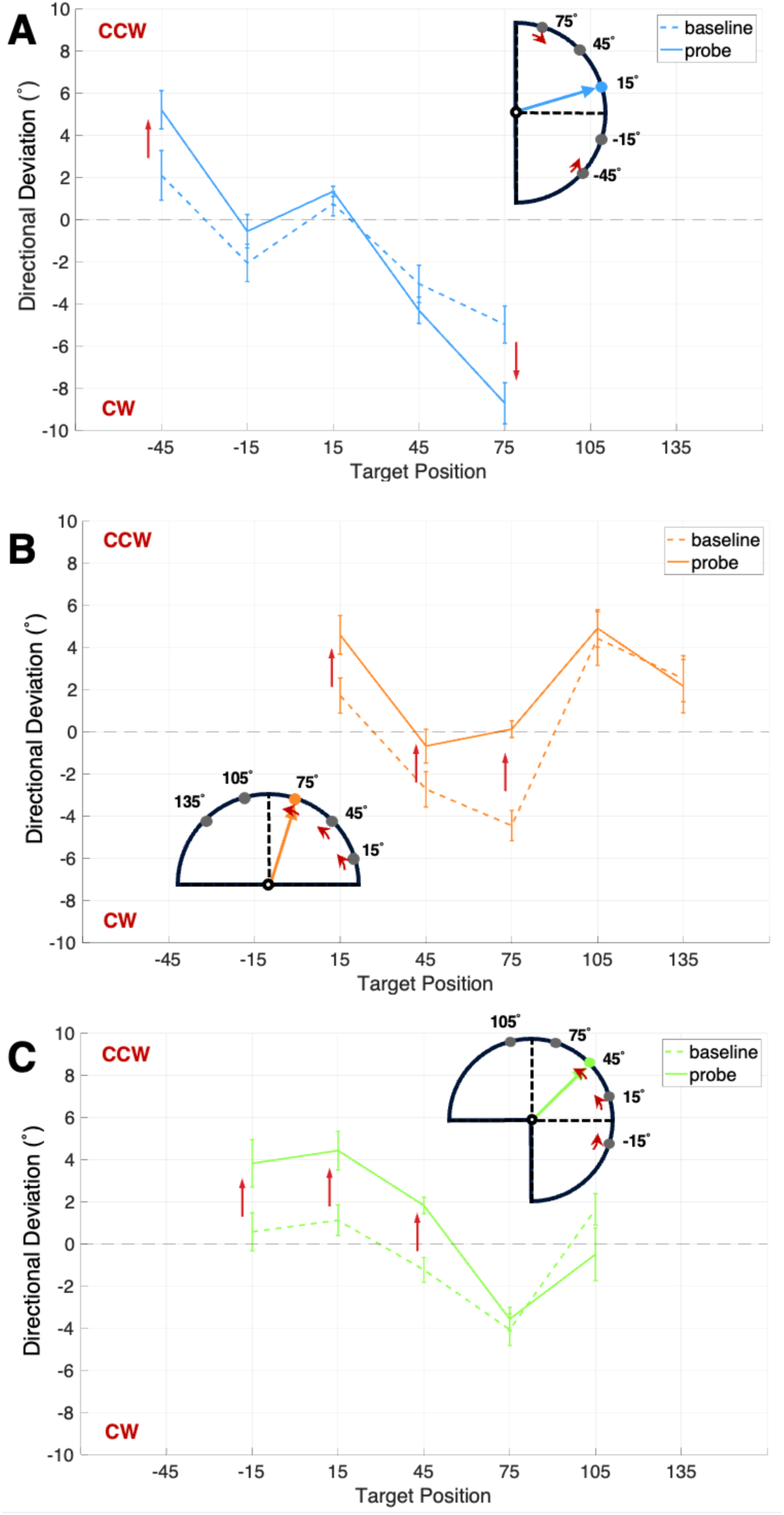
Average reaching error during baseline and probe trials for three groups at each target position. Dashed lines represent baseline trials and solid lines represent probe trials. Red arrows indicate the direction of bias changes that reached significance or marginally significance, demonstrating the effect of repetition. All significant deviations at non-repeated targets were directed toward the repeated target direction, confirming the attractive nature of use-dependent biases.

Similarly, both the Repelling and Control groups demonstrated significant counterclockwise deviations at probes positioned clockwise to their respective repetition directions. As reported in section 3.1, participants exhibited substantial intrinsic biases at certain directions (e.g., −2.33° ± 0.45° at the 45° target and −4.48° ± 0.44° at the 75° target). In contrast, UDL in reaching tasks typically induces relatively modest biases; the largest bias observed across all directions and conditions was only 3.74° ± 0.61° (at the 75° target in the Attracting group). Given this disparity in magnitude, UDL effects may not have reached statistical significance at all probe locations, potentially being obscured by the larger intrinsic biases. Critically, however, all statistically significant deviations were directed toward the respective repetition target, establishing a clear attractive bias pattern. We observed no repulsive effects at any target location.

### Use-dependent biases affect the initial state of error-based adaptation but not learning dynamics

Having established that repetitive training induced use-dependent biases, we next examined whether these biases influenced subsequent implicit EBL. Following the repetition phase, all three groups transitioned to an EC task where they reached the 45° target while the cursor was constrained to move along 75° regardless of actual hand direction. This paradigm primarily engages implicit adaptation processes while minimizing explicit strategic responses (Morehead et al., 2017).

All groups demonstrated immediate adaptive responses when the visual perturbation was introduced, exhibiting clockwise changes in hand direction to compensate for the counterclockwise clamped cursor (Figure 5A). However, their initial trial clearly reflected the UDL bias: the Attracting group showed substantial deviation, relative to the target, toward their previous repetition direction (8.61° ± 1.06°) — a clockwise bias that was in the to-be-adapted direction. The Repelling group exhibited a counterclockwise bias (−2.29° ± 1.67°), also directed toward their repetition target but opposing the required adaptation. The Control group exhibited a small deviation, close to zero (1.72° ± 0.56°). The difference among the three groups was significant (*F*(2,53) = 22.320, *p* < .001), and pairwise comparisons further indicated that all three groups differed significantly with each other (Attracting vs. Repelling: *t* = 6.576, *p* < .001; Attracting vs. Control: *t* = 4.157, *p* < .001; Repelling vs. Control: *t* = 2.357, *p* = .022).

**Figure 5.**
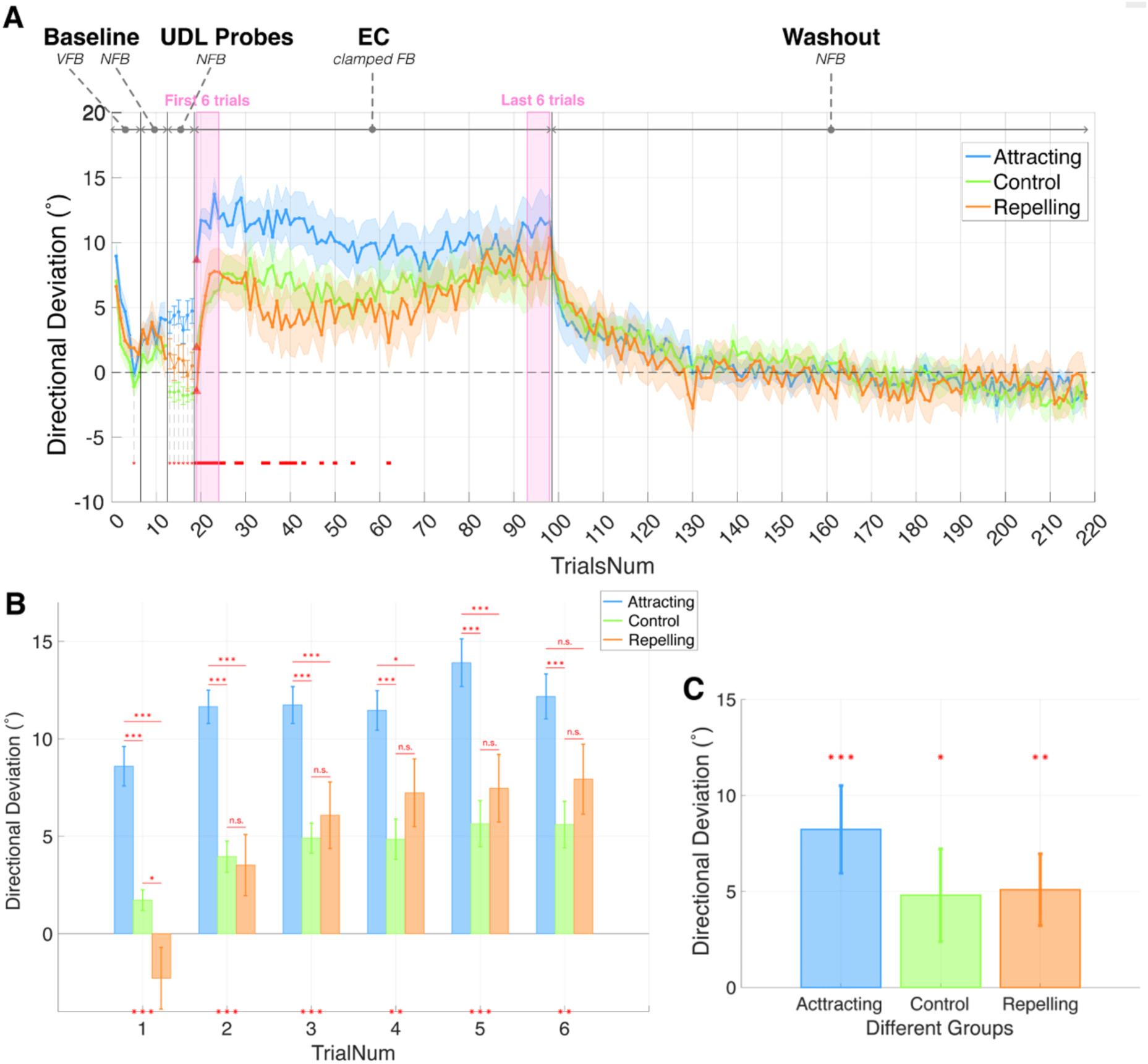
Main results of the process of EC adaptation. **(A)** Hand deviation trajectories for the 45° target from baseline through clamp phases. Shaded areas represent standard errors. Vertical gray lines delineate task phases: baseline with veridical feedback (VFB), baseline without feedback (NFB), probe trials without feedback, EC task with clamped feedback, and washout phase without feedback. The red bar in the EC phase indicates trials with significant one-way ANOVA results (p < .05). Following repetition training, the three groups exhibited separated reaching endpoints. Upon entering the EC task, all groups rapidly adapted to the clamped error, ultimately achieving comparable adaptation levels. **(B)** Reaching error during the first six EC trials. In the first trial, all three groups differed significantly from each other. From the second trial onward, the Repelling and Control groups converged, showing substantially smaller biases than the Attracting group. **(C)** Adaptation levels during the EC task, calculated as the difference between final EC trials and baseline. All three groups achieved final adaptation extents that substantially exceeded baseline levels, confirming successful engagement of implicit EBL. No significant differences were observed between groups.

Although overall group difference persists in the following trials (e.g., 2^nd^: *F*(2,57) = 16.350, *p* < .001; 6^th^: *F*(2,57) = 5.017, *p* = .010), we found that the difference between the Repelling and Control groups diminished rapidly, becoming non-significant by the second trial (Figure 5B; post-hoc comparison between Repelling and Control groups for the 2^nd^ trial: *t* = 0.271, *p* = .787). In subsequent trials, these two groups did not differ anymore (Figure 5B). The large group difference between the Attracting group and the other two groups diminished more slowly, but all groups converged by the final adaptation trials (last 6 trials; *F*(2,57) = 0.745, *p* = .479). Thus, after extended EC learning, all groups achieved significant and comparable levels of implicit adaptation (Figure 5C; Attracting: 8.23° ± 2.28°, *t(19)* = 3.611, *p* < .001; Repelling: 5.09° ± 1.87°, *t(19)* = 2.720, *p* = .007; Control: 4.81° ± 2.41°, *t(19)* = 1.992, *p* = .030), despite their initial difference caused by UDL.

To further examine whether UDL influenced the dynamics of EC adaptation, we analyzed learning rates quantified as trial-to-trial changes in reaching angle throughout the EC phase. If UDL altered error sensitivity of implicit adaptation, we would expect systematic differences in how rapidly participants adjusted their movements in response to clamped feedback. However, we observed no significant group differences throughout the adaptation phase (Figure 6). This absence of group differences extended across all learning trials, even when analyzing trial-by-trial changes after the critical first trial (*F*(2,57) = 2.935, *p* = .061).

**Figure 6.**
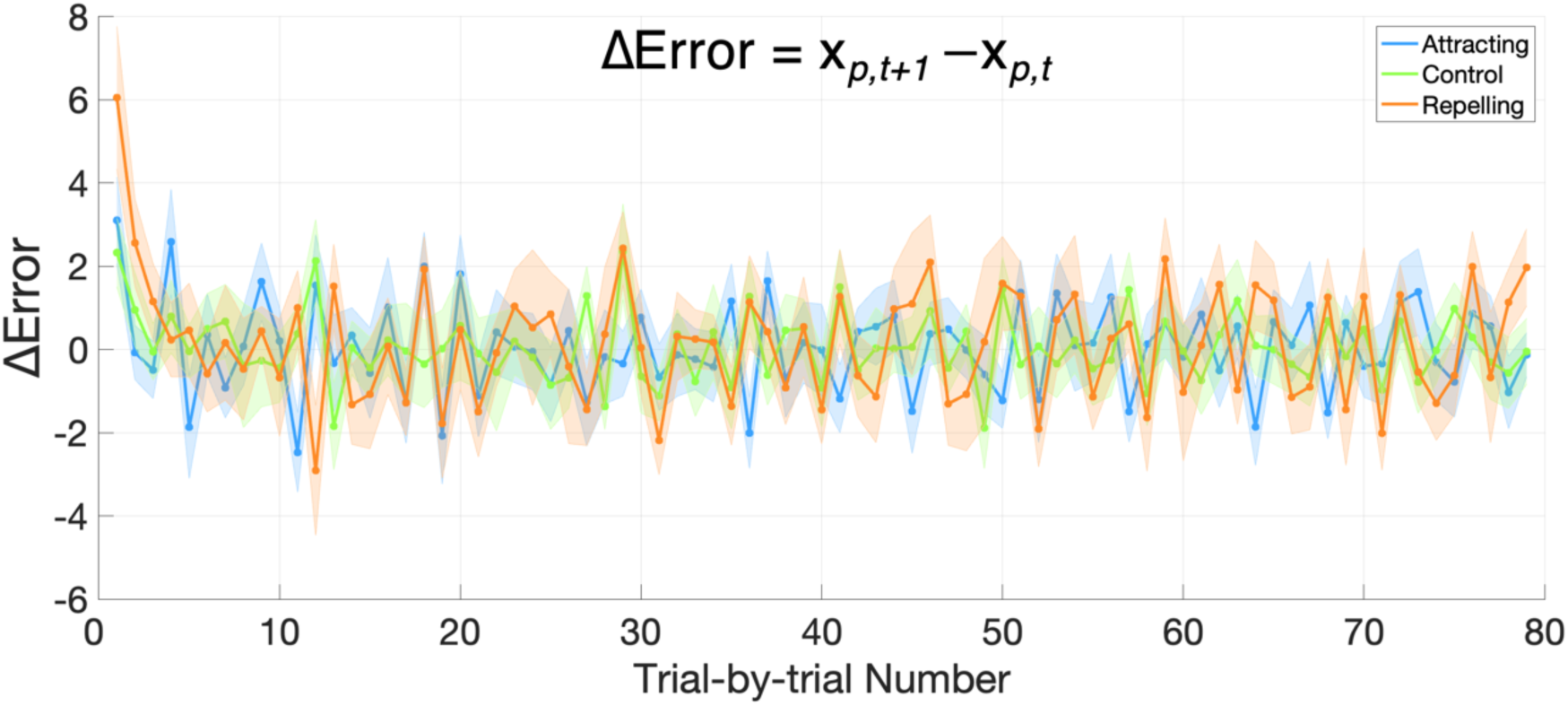
Trial-by-trial changes in reaching angle during the EC task. No significant differences were observed between groups at any point, indicating that UDL does not modulate the rate of implicit EBL.

### The temporal dynamics of use-dependent biases during and after EC adaptation

The preceding analyses revealed that use-dependent effects were confined to early adaptation trials, with their influence diminishing progressively over time. To elucidate this temporal evolution, we normalized the Attracting and Repelling groups’ performance by subtracting the Control group’s performance, thereby isolating the contribution of UDL during EC learning (Figure 7). Indeed, the UDL exerted large initial influence: both conditions showed normalized deviation toward their respective repetition directions, clockwise for Attracting (6.89° ± 1.06°) and counterclockwise for Repelling (−3.78° ± 1.51°) groups. As the adaptation progressed, the group difference remained significant for approximately 30 trials but decreased gradually. In the average of the final 6 trials, normalized errors were 5.24° ± 2.02° for the Attracting group and 1.79° ± 2.04° for the Repelling group, with no significant difference between them (*t(38)* = 1.205, *p* = .236). This result echoes the comparable final adaptation levels observed without normalization (Figure 5C). And the gradual decay pattern of UDL bias was consistent with previous reports (e.g., Diedrichsen et al., 2010). Though both groups gradually reduced their UDL influences, their decay patterns differed, with the Attracting group decaying more slowly than the Repelling group. This asymmetry suggests that persistence of the UDL bias varies across target directions.

**Figure 7.**
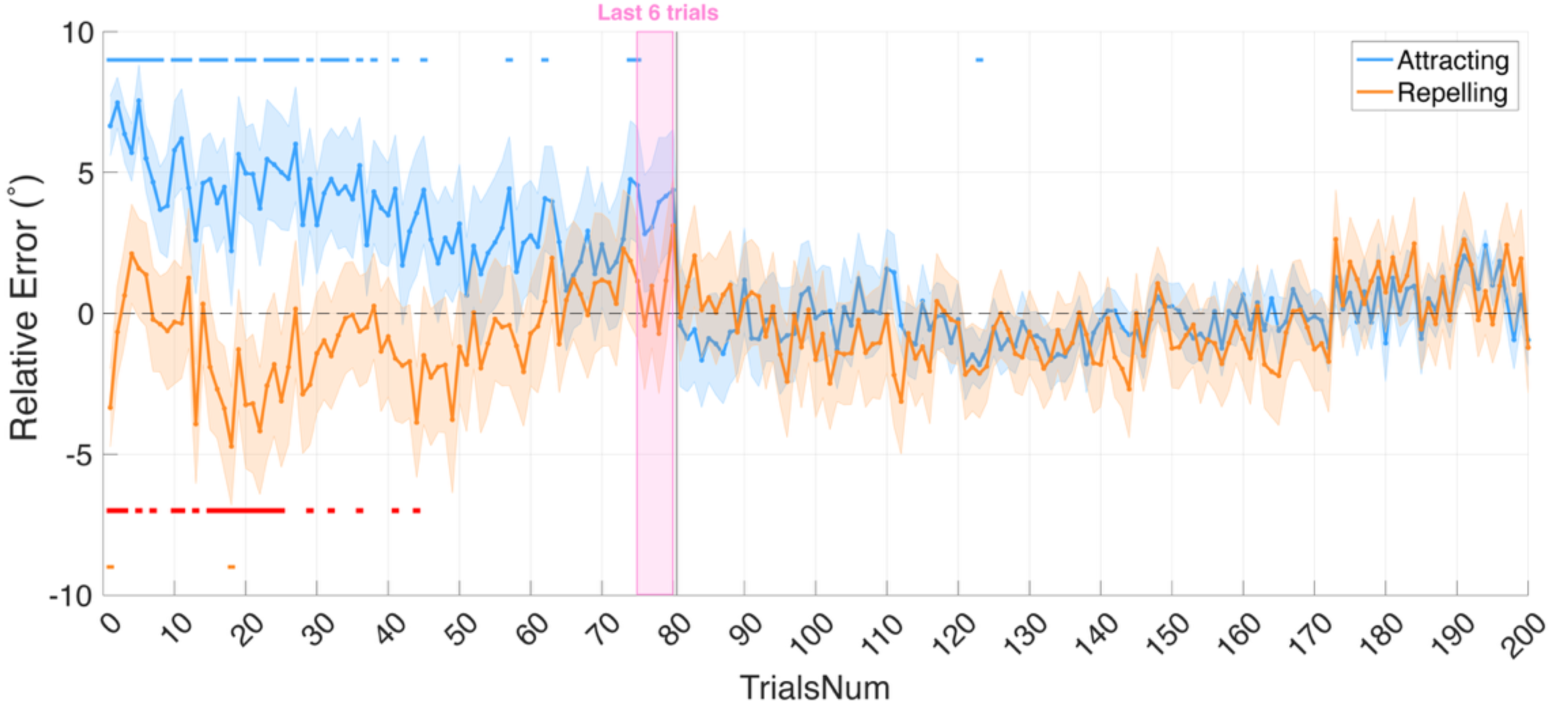
Time course of the UDL effect, isolated by subtracting the Control condition. The difference of reaching biases toward the repeated direction between the two groups reached maximum at the first EC trial. Subsequently, both curves decayed gradually but with different temporal dynamics: the Repelling group showed rapid decay toward zero, while the Attracting group demonstrated slower decay and larger effect magnitudes throughout the adaptation phase. The red bar indicates trials with significant one-way ANOVA results (p < .05), while the blue and orange bars indicate significant differences from zero for the Attracting group and the Repelling groups, respectively.

## Discussion

The present study investigated whether use-dependent learning (UDL), a form of motor plasticity induced by repetitive movement, influences subsequent implicit error-based learning (EBL). Our participants first performed extensive repetitive reaching to establish use-dependent biases, then immediately transitioned to an error-clamp (EC) adaptation task that selectively induced implicit learning processes. We manipulated the spatial relationship between repeated and adapted directions across three groups: Attracting (UDL bias aligned with the to-be-adapted direction), Repelling (UDL bias opposite the to-be-adapted direction), and Control (no directional conflict). Our results demonstrate that repetitive reaching successfully induces use-dependent biases, attracting subsequent movements toward the repeated direction. These biases substantially affected movement directions during initial adaptation, with the three groups exhibiting markedly different starting points for the subsequent implicit error-based adaptation. However, UDL did not alter adaptation dynamics—all groups exhibited comparable learning rates and converged to similar asymptotic levels despite their distinct initial states. These findings support an independent rather than interactive relationship: UDL biases shift the initial state of adaptation but do not affect the implicit EBL process itself.

Our study addressed whether UDL bias merely shifts the starting state for adaptation (independent hypothesis) or fundamentally alters the learning dynamics of implicit EBL (interaction hypothesis). Our findings strongly favor their independence. Although the three participant groups began EC adaptation with markedly different initial deviations reflecting their UDL histories, they showed no differences in trial-by-trial learning rates or final adaptation levels. The Attracting group, despite beginning with substantial bias toward the to-be-adapted direction, achieved no greater asymptotic adaptation than controls. Similarly, the Repelling group showed no reduced final adaptation despite opposing initial bias. Trial-by-trial changes in reaching angle also revealed no systematic group differences throughout adaptation. These findings indicate that use-dependent bias acts as an offset in the initial movement plan, but once EC adaptation begins, implicit learning responds to motor errors with unchanged sensitivity.

These findings extend previous investigations of UDL-EBL relationships. Diedrichsen et al. (2010)used a force field to induce a UDL bias and found that subsequent EBL could correct these directional biases without visual errors, presumably relying on proprioceptive (Shadmehr & Mussa-Ivaldi, 1994) or perceptual error in hand localization (Zhang et al., 2024). However, they did not address whether one process influences the other. In contrast, our study directly addresses the interaction problem by manipulating UDL bias in either the facilitating or opposing direction of the subsequent EBL. Huang et al. (2011) also found that hand movement repetition during visuomotor rotation produced a robust directional bias in initial movement trajectories. However, visuomotor rotation learning includes substantial explicit components that shape overall learning rates. Thus, whether implicit learning processes are affected by UDL bias remains unresolved by their investigation. By employing the EC paradigm, we isolated implicit adaptation and provided direct evidence that UDL and implicit EBL operate independently.

More broadly, our findings contribute to understanding how multiple learning systems coordinate during motor learning. Categorized by learning signals, motor learning arises from three distinct mechanisms: reinforcement learning (RL) driven by reward, UDL driven by repetition, and EBL driven by sensory prediction error (Wolpert et al., 2011). Previous research focused predominantly on interactions between RL and EBL during motor adaptation, demonstrating that reward signals modulate error-based adaptation through metacognitive or motivational gating (Galea et al., 2015; Kim et al., 2019; Sugiyama et al., 2023). In contrast, our study examined two implicit—implicit EBL and UDL—which have both been viewed as changes in motor execution (Morehead et al., 2017; Tsay et al., 2022), and found no evidence for interaction.

This pattern suggests that the architecture of motor learning may be hierarchically organized: whereas RL represents a higher-level cognitive process that can flexibly gate or weight lower-level modules, UDL and implicit EBL appear to function as parallel modules that contribute additively to motor output. We argue that maintaining independence among execution-level mechanisms offers a balance between stability and flexibility: UDL stabilizes motor output by reinforcing recently executed actions, while implicit EBL recalibrates the sensorimotor system based on prediction errors to accommodate possible environmental and bodily changes.

We postulate that whether different learning mechanisms show coordinated changes, sometimes appearing to interact, depends on whether they share common learning signals or overlapping neural substrates. For common signals, interactions may emerge when mechanisms use compatible error information. For example, RL and EBL jointly contribute to performance when both processes use the magnitude of target error as a teaching signal in a visuomotor rotation task (Izawa & Shadmehr, 2011). In contrast, UDL is driven by Hebbian strengthening of repeated motor commands (Verstynen & Sabes, 2011), while implicit EBL mainly depends on prediction errors (e.g., Mazzoni & Krakauer, 2006; Taylor & Ivry, 2011; Tseng et al., 2007). Additionally, interaction may depend on neural substrate overlaps. Torrecillos et al. (2014) showed that sensory-prediction errors and reward-prediction errors evoke highly similar frontocentral ERPs, with shared generators in pre-SMA and surrounding medial frontal areas, indicating a neural overlap between RL and EBL. However, UDL predominantly involves Hebbian-like plasticity in primary motor cortex (M1) and premotor areas (Classen et al., 1998; Verstynen & Sabes, 2011), whereas implicit EBL relies primarily on cerebellar circuits (Galea et al., 2011; Schlerf et al., 2012). Although the cerebellum and M1 are interconnected, their functional interaction appears task-specific. During EBL, cerebellar output modulates M1 excitability (Jayaram et al., 2011). During UDL, however, neural evidence shows changes localized to M1 without any evidence of cerebellar excitability changes (Spampinato & Celnik, 2021). Our behavioral findings echo this neural dissociation, suggesting that M1-driven use-dependent plasticity operates independently of cerebellar error-processing mechanisms during implicit learning.

Although we observed an overall effect of use-dependent bias among all the three groups, the UDL effect was not universally observed for all the probe targets. This heterogeneity may reflect several factors. First, previous studies typically averaged the UDL effect across targets that are symmetrically positioned relative to the repeated target (Marinovic et al., 2017; Tsay et al., 2022; Verstynen & Sabes, 2011). When we also used this type of data collapse, our results became more consistent and in fact resembled their findings: both angular positions showed significant bias toward the repeated direction (30° deviation: 1.19° ± 0.40°, *t*(59) = 2.968, *p* = .004; 60° deviation: 2.58° ± 0.44°, *t*(59) = 5.805, *p* < .001), with larger biases at positions farther away from the repeated target (*t*(59) = 4.175, *p* < .001). Another factor that contributes to inconsistent UDL bias across targets is that people showed different baseline movement biases across targets, before any UDL training (Figure 3). This finding was also revealed by a recent large-sample study (Wang et al., 2024), and we believe these baseline biases could obscure UDL effects in some directions.

The EC adaptation extent observed in our experiment was smaller than typically reported in some studies (e.g., Bond & Taylor, 2015; Morehead et al., 2017; Zhang et al., 2024). This attenuation of implicit adaptation is likely caused by our extended baseline with veridical feedback. A recent study found a similar reduction in implicit adaptation after a long baseline. They argued that prolonged baseline training strengthens prior expectations about feedback consistency, which in turn reduces the extent to which the motor system incorporates EC feedback (Avraham & Ivry, 2024). Nevertheless, the fact that all three groups of participants converged to a similar, albeit smaller, final adaptation extent supports the lack of interaction between UDL and implicit EBL.

We also observed an asymmetry in the persistence of UDL bias between the Attracting and Repelling groups: the Attracting group’s bias decayed more slowly than the Repelling group’s. This phenomenon has not been addressed previously and suggests that the effect of UDL bias may vary across angular locations, with stronger or more persistent effects at certain directions. Of note, we examined only one temporal ordering of learning conditions, i.e., UDL followed by implicit EBL. Future studies should examine alternative sequences or simultaneous engagement of both mechanisms to test whether their relationship will persist across different learning contexts.

## Conclusions

This study provides evidence that UDL and implicit EBL combine additively during motor adaptation. Repetition-induced biases substantially affect initial adaptation states but do not modulate the EBL process itself: all groups converged to similar asymptotic levels at comparable rates despite their distinct initial learning states induced by UDL. These findings advance our understanding of how multiple implicit learning systems coordinate during motor behavior and suggest that “low-level” execution-related motor learning components are additive. Future research should examine whether this additive relationship generalizes across different temporal sequences and spatial configurations.

## Data availability

Data are available at: https://osf.io/mk538/

## Author contributions

YL: Conceptualization, Methodology, Software, Data Curation, Formal Analysis, Visualization, Writing; KW: Conceptualization, Methodology, Supervision, Funding Acquisition, Writing

